# Cell type-specific variation of somatotopic precision across corticostriatal projections

**DOI:** 10.1101/261446

**Authors:** Bryan M. Hooks, Andrew E. Papale, Ronald Paletzki, Muhammad Feroze, Brian S. Eastwood, Jonathan J. Couey, Johan Winnubst, Jayaram Chandrashekar, Charles R. Gerfen

## Abstract

The striatum shows general topographic organization and regional differences in behavioral functions. How corticostriatal topography differs across cortical areas and cell types to support these distinct functions is unclear. This study contrasted corticostriatal projections from two layer 5 cell types, intratelencephalic (IT-type) and pyramidal tract (PT-type) neurons, using viral vectors expressing fluorescent reporters in Cre-driver mice. Long-range corticostriatal projections from sensory and motor cortex are somatotopic, with a decreasing somatotopic specificity as injections move from sensory to motor and frontal areas. Somatotopic organization differs between IT-type and PT-type neurons, including injections in the same site, with IT-type neurons having higher somatotopic stereotypy than PT-type neurons. Furthermore, IT-type projections from interconnected cortical areas have stronger correlations in corticostriatal targeting than PT-type projections do. Thus, as predicted by a long-standing basal ganglia model, corticostriatal projections of interconnected cortical areas form parallel circuits in basal ganglia-thalamus-cortex loops.

## Introduction

Primary motor (M1) and primary somatosensory (S1) areas of cerebral cortex are somatotopically organized, with distinct body regions represented in adjacent areas. Though sensory and motor cortices specialize in distinct functions, corticocortical projections reciprocally connect them. Similarly, corticostriatal inputs are topographically organized. Overlaid on this pattern, however, output from any given cortical area projects broadly and overlaps with output from other areas, including topographically related ones^1,2^. A longstanding model of corticostriatal organization is that striatal regions integrate input from multiple cortical areas that are functionally interconnected^3,4^. This suggests that the striatum is organized into distinct regions^2^ associated with different behavioral functions^5,6^. While there is topographic organization, different functions of dorsolateral, dorsomedial, and ventral divisions are not strictly topographic^7,8^. To better understand how information from the cortex is integrated within the striatum, this study first asks whether projections from different cortical areas project to stereotyped somatotopic sectors of striatum across animals by quantifying overlap and segregation between sensory, motor, and frontal projections. As a subsequent step, this data tests whether corticocortical connectivity predicts convergence or interdigitation within the striatum.

Addressing these questions is not straightforward with conventional anatomical techniques, since the corticostriatal projection originates from two distinct excitatory neuron categories in layer 5 (L5): pyramidal tract type (PT-type) neurons and intratelencephalic (IT-type) neurons^9,10^. PT-type neurons send projections to the thalamus, subthalamic nucleus, superior colliculus and brainstem with collaterals in ipsilateral striatum^11^, but do not project to contralateral cortex nor contralateral striatum. In contrast, IT-type cells project exclusively to ipsi- and contralateral striatum and cortex, and not to subcortical targets^10^. In motor areas, local circuits are hierarchically organized such that IT-type cells connect to each other and project to PT-type neurons, but PT-type neurons do not connect to IT-type cells^12^. Thus, information at different stages of processing is transmitted out of cortex, conveying distinct messages^13^.

The differences between the corticostriatal projections of these two major cell types were analyzed using stereotaxic injection of Cre-dependent reporters into sensory, motor, and frontal cortical areas of Cre-driver mice selective for IT-type and PT-type neurons. Sectioned brains were then imaged and aligned to a reference brain, the Mouse Common Coordinate Framework version 3 (CCF v3)^14-16^ to quantify axonal fluorescence in a standard coordinate system. Targeting of axonal projections in striatum and other targets of motor and sensory output was quantified to assess the somatotopic organization of projections. This data reveals that the somatotopic organization of projections differs between IT-type and PT-type neurons and between sensory and motor areas. Thus, the information cortex provides for striatal processing differs across these two cortical output channels.

## Results

### Generation of a dense library of IT-type and PT-type corticostriatal projections

To analyze the corticostriatal projections of specific pyramidal cell types, mouse lines selectively expressing Cre in IT-type (Tlx3_PL56) and PT-type (Sim1_KJ18) neurons^17^ were injected with AAV expressing Cre-dependent tracers. Each mouse received injections of 3 different AAV vectors (GFP, td-tomato, and smFPs; Table 1^18^) into different locations of sensory, motor and frontal cortex (Fig. 1 and Supplementary Fig. 1). A whole-brain reconstruction from tiled images^19^ (Supplementary Fig. 1b-e) was registered to a common reference frame using BrainMaker software (MBF Bioscience) with alignment precision of ∼50-70 μm (Supplementary Fig. 1l-y). Original images were posted at: http://gerfenc.biolucida.net/link?l=Jl1tV7. Placing all voxels from all brains in the same reference space enabled quantitative analysis of regions of interest across different animals (Supplementary Fig. 1h-i).

**Figure 1.**
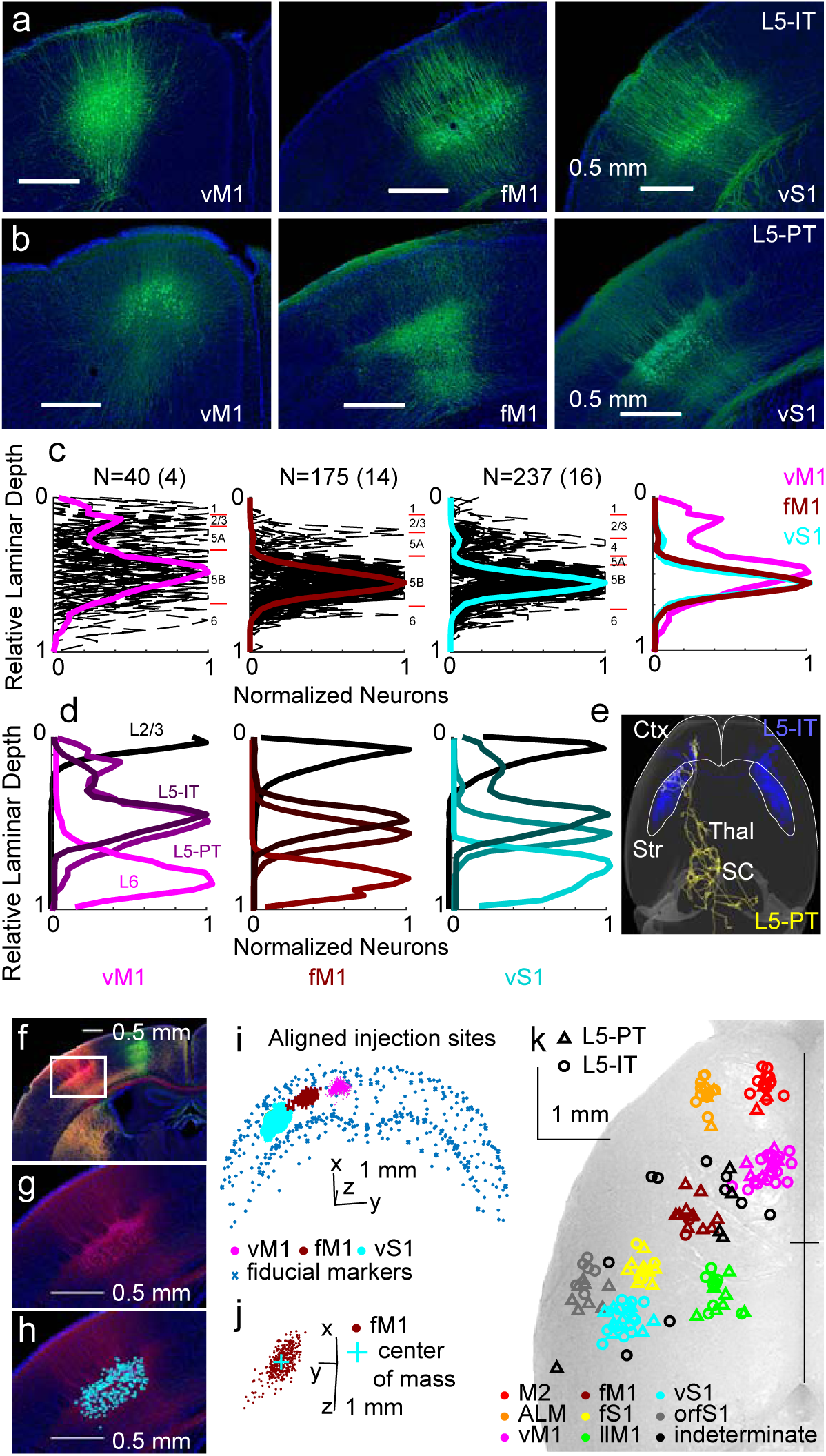
Cre-driver lines label specific pyramidal neuron cell types. (a, b) Example coronal images at the injection site of Tlx3_PL56 (a, labeled L5-IT) and Sim1_KJ18 (b, labeled L5-PT) in vM1, fM1, and S1 to show soma location. Scale bars, 0.5 mm. All images pseudocolored green for comparison. (c) Quantification of soma location of Sim1_KJ18 mice injected in vM1, fM1, and S1. Comparison across regions shown at right. N indicated shows # of sections (# of mice) quantified. Purple, vM1; burgundy, fM1; teal, vS1. Pia is at relative laminar depth of 0; white matter is at 1. Black dashed lines represent individual sections with each section normalized to 1. Red tick marks show estimated laminar borders for cortical layers. (d) Mean neuron distribution for four lines labeling L2/3 (Sepw1_NP39), L5-IT (Tlx3_PL56), L5-PT (Sim1_KJ18), and L6 (Ntsr1_GN220) in three cortical areas. (e) Different targets of IT-type (Tlx3_PL56) and PT-type (Sim1_KJ18) neurons, illustrated with single axon reconstructions: IT-type neurons (blue) project to ipsi- and contralateral cortex (Ctx) and striatum (Str), while PT-type neurons (gold) target ipsilateral cortex and striatum, as well as subcortical targets in thalamus (Thal), superior colliculus (SC) and brainstem. (f-h) Low and high magnification images in Neurolucida of an injection site in a Sim1_KJ18 mouse. White box in (f) indicates magnified area (g and h). Scale bars 0.5 mm. (h) Annotation of somata at injection site in Neurolucida. Blue circles (for red injection) indicate AAV-infected cell bodies expressing smFPs. (i and j) Coordinates of somata from Neurolucida and fiducial markers placed along the pial surface of cortex and white matter were aligned to the CCF using the same coordinate transform as for the structure channel of the given brain. The somata for three injections (teal, burgundy, and purple) and fiducial markers (gray) were then plotted in 3-d (axes as indicated, with 1 mm scale bar, coronal viewpoint). This projection was rotated (j) for a dorsal view showing the center of mass (teal) for the burgundy injection and the anterior/posterior spread of infected somata. (k) Center of mass of Tlx3_PL56 (N%92, circles) and Sim1_KJ18 (N% 62, triangles) injections plotted in the CCF and spatially clustered. Eight clusters are shown in red (M2), orange (ALM), purple (vM1), burgundy (fM1), green (llM1), yellow (fS1), teal (vS1), and gray (orfS1). Indeterminate injection sites are in black. Sites are superimposed on an image of the dorsal surface of mouse cortex. Black cross marks midline and bregma.

**Table 1.**
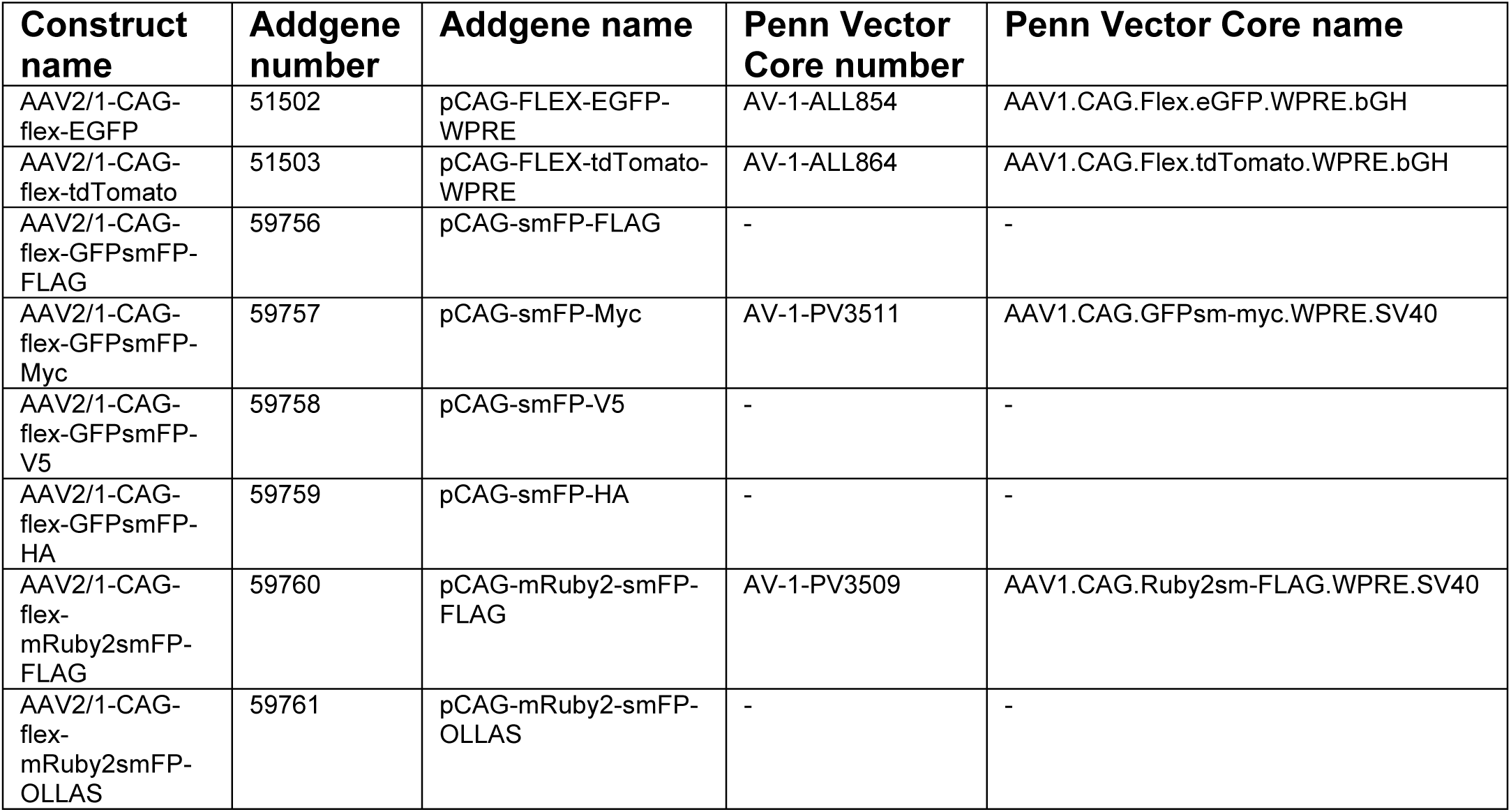
**Constructs for Tracing (Online Methods)**

| **Construct name** | **Addgene number** | **Addgene name** | **Penn Vector Core number** | **Penn Vector Core name** |
| --- | --- | --- | --- | --- |
| AAV2/1-CAG-flex-EGFP | 51502 | pCAG-FLEX-EGFP-WPRE | AV-1-ALL854 | AAV1.CAG.Flex.eGFP.WPRE.bGH |
| AAV2/1-CAG-flex-tdTomato | 51503 | pCAG-FLEX-tdTomato-WPRE | AV-1-ALL864 | AAV1.CAG.Flex.tdTomato.WPRE.bGH |
| AAV2/1-CAG-flex-GFPsmFP-FLAG | 59756 | pCAG-smFP-FLAG | - | - |
| AAV2/1-CAG-flex-GFPsmFP-Myc | 59757 | pCAG-smFP-Myc | AV-1-PV3511 | AAV1.CAG.GFPsm-myc.WPRE.SV40 |
| AAV2/1-CAG-flex-GFPsmFP-V5 | 59758 | pCAG-smFP-V5 | - | - |
| AAV2/1-CAG-flex-GFPsmFP-HA | 59759 | pCAG-smFP-HA | - | - |
| AAV2/1-CAG-flex-mRuby2smFP-FLAG | 59760 | pCAG-mRuby2-smFP-FLAG | AV-1-PV3509 | AAV1.CAG.Ruby2sm-FLAG.WPRE.SV40 |
| AAV2/1-CAG-flex-mRuby2smFP-OLLAS | 59761 | pCAG-mRuby2-smFP-OLLAS | - | - |

As expected for IT-type neurons, injections in Tlx3_PL56 mice labeled axonal projections that bilaterally targeted cortex and striatum, but not other subcortical structures^10^ (Fig. 1e). By contrast, axonal projections in the Sim1_KJ18 line were restricted to the hemisphere ipsilateral to the injection within the cortex and striatum. Labeled neurons also projected to the thalamus, subthalamic nucleus, superior colliculus, pontine and medullary nuclei, typical of PT-type corticofugal neurons^11^. IT-type neurons are generally located in more superficial layer 5 than PT-type neurons, with considerable overlap. Injections in Sim1_KJ18 and Tlx3_PL56 infected a small number of L2/3 neurons. Somata of labeled pyramidal neurons at injection sites were marked in Neurolucida and their relative laminar depth plotted (Fig. 1a-d). Tlx3_PL56 and Sim1_KJ18 labeled neurons at injection sites were consistent with prior descriptions of the laminar locations of IT and PT neurons^20,21^.

The coordinates of labeled somata for each injection in the original images were marked and transformed into the CCF reference frame (Fig. 1f-k), with the average used to determine a center of mass for the injection site (Fig. 1j). The center of mass was used to cluster injection sites for Sim1_KJ18 and Tlx3_PL56 into 8 clusters across sensory, motor and frontal cortex (Fig. 1k). These corresponded to vibrissal, forelimb, and orofacial somatosensory cortices (vS1, fS1, and orfS1); vibrissal, forelimb, and lower limb motor cortices (vM1, fM1, and llM1); and frontal areas (anterior lateral motor cortex (ALM) and secondary motor cortex (M2)). Indeterminate injection sites (black) were not clustered. The names assigned to these sites correspond to microstimulation mapping for motor areas^22,23^ and somatotopic mapping of sensory areas^24-26^.

A methodology was developed to quantitatively compare projections from different injections sites. Images were thresholded to eliminate 99% of background (Supplementary Fig. 1z). Three example injection sites (from Tlx3_PL56 mice in vM1, vS1, and ALM) illustrate the methodology for comparison (Fig. 2). Suprathreshold voxel intensity for ipsilateral striatum was compared on a voxel-by-voxel basis using voxels that were suprathreshold for *both* channels (Fig. 2a). The Pearson correlation coefficient (PCC) was used to assess the relationship within the striatum for each pair of injections (Fig. 2b). To localize where within the striatum correlations occurred, correlation was computed for each plane along the anterior/posterior axis (Fig. 2c-e). Correlation values varied dependent on both the particular injection sites and the rostro-caudal level of the striatum. In the example shown, correlation was near zero in anterior striatum, but became well correlated for vS1 and vM1 in mid- and posterior ipsilateral striatum (black line). In contrast, correlation is negative for both vS1 and vM1 when compared to the ALM injection (yellow and blue lines, Fig. 2e-f). Correlations were noisier when measured based on small numbers of voxels (anterior and posterior poles of striatum, Fig. 2e-f). The general pattern was similar for individual injections (Fig. 2e) compared to the population (Fig. 2f), but the magnitude of correlation varied considerably depending on individual M1 and S1 injections considered. This anatomical overlap of afferents corresponds to shared targeting of functional synaptic output to specific single neurons. This was tested using a dual channel circuit mapping approach with conventional ChR2 and red-shifted ReaChR^27^ expressed in vM1 and vS1 respectively. Whole cell recordings from striatal projection neurons (SPNs) in the overlapping region of vM1 and vS1 projections revealed synaptic convergence in all neurons recorded (Supplementary Fig. 2). This confirmed that convergent axonal projections, such as those from somatotopically aligned regions of sensory and motor cortex, also shared functional synaptic targets.

**Figure 2.**
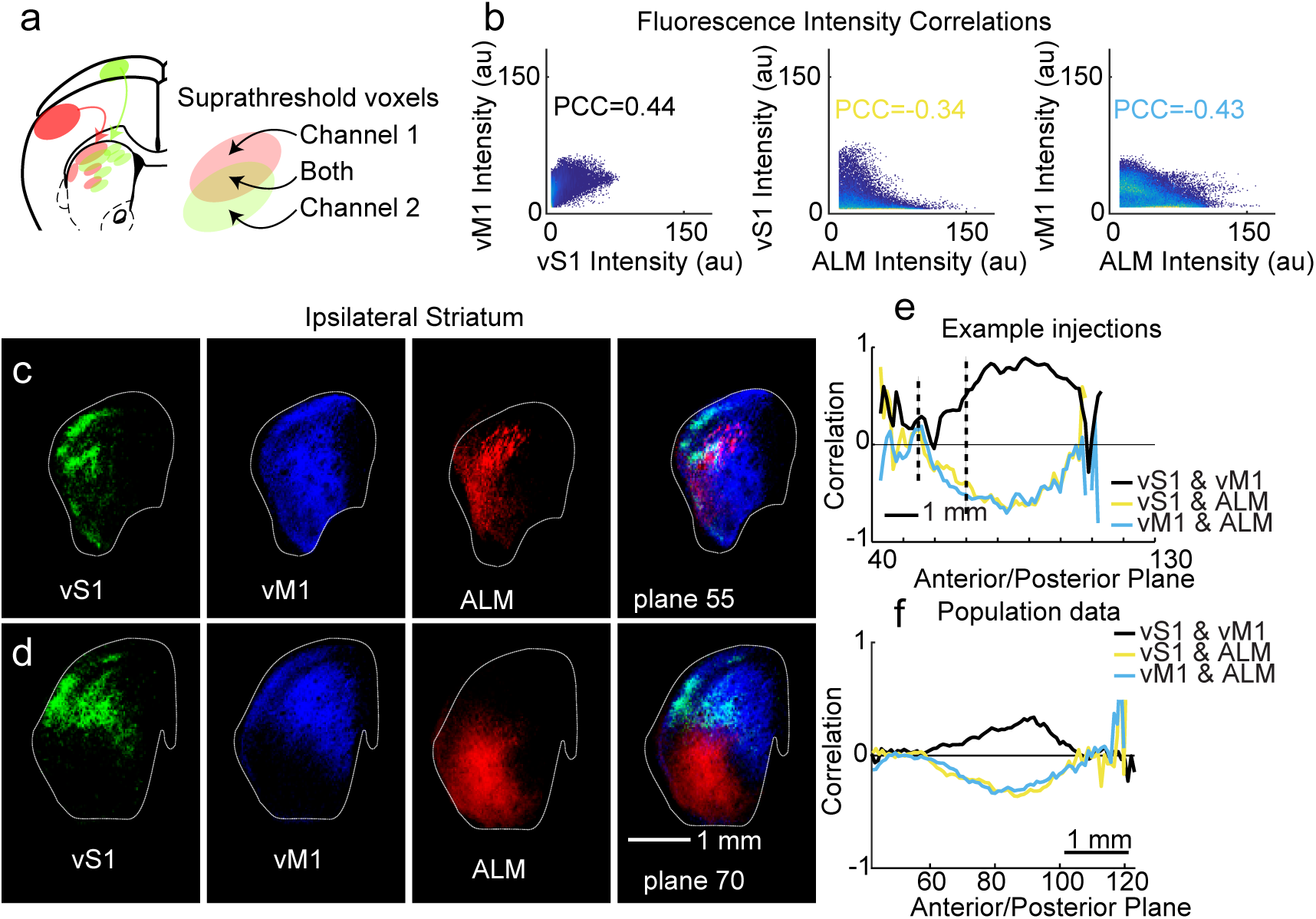
Computation of correlation for projections to ipsilateral striatum. (a) Correlation for two injections (red and green channels) in a given structure (striatum, illustrated) is computed based on voxels where both channels are suprathreshold. (b) For three example Tlx3_PL56 (IT-type) mouse injections in vM1, vS1, and ALM, scatterplot of all voxel intensities (8-bit imaging; arbitrary units) in ipsilateral striatum for two injections. Blank space between axis and points indicates threshold. Individual points in dark blue; multiple points increase yellow intensity. (c-d) Example coronal images from aligned brains in corresponding planes showing arborization of IT-type axons in ipsilateral striatum. vS1 shown in green, vM1 in blue, and ALM in red. (e) Correlation coefficient as a function of anterior/posterior plane in ipsilateral striatum for pairwise comparisons between three example injections. Correlation is noisy at anterior and posterior poles of striatum due to small voxel numbers in those planes. (f) Population mean correlation coefficient as a function of anterior/posterior plane in ipsilateral striatum for pairwise comparisons (the mean correlation for each vS1 compared to each vM1 in black, for example). vS1 and vM1 comparison, N=340 injection pairs; vS1 and ALM comparison, N=204; vM1 and ALM comparison, N=240.

### Sensory, motor, and frontal corticostriatal projections target somatotopically specific areas

To study somatotopy of ipsilateral corticostriatal projections, this analysis was extrapolated to all eight injection clusters, which included sensory areas (vS1, fS1, and orfS1), motor areas (vM1, fM1, and llM1), and frontal areas (ALM and M2). Sensory, motor, and frontal areas were taken to be three modalities for cortical function, with the clusters within each modality representing different somatotopic regions (whisker, forelimb, and hindlimb for example) within that modality. Projections from different parts of the same cortical modality displayed a topographic organization along the rostral to caudal axis, demonstrated by the relationship of the projection of the aforementioned sensory, motor, and frontal areas (Fig. 3a). This demonstrated the maintenance of the somatotopic organization within modalities in their projections to the striatum. On the other hand, comparison of the projections between sensory, motor and frontal areas showed considerable overlap (Fig. 3b). Quantitative analysis reveals varying levels of input from cortical areas along the rostro-caudal axis (Fig. 3c). Somatosensory injections were biased towards more posterior sites, with maximum intensity and suprathreshold voxel numbers peaking more caudally than motor or frontal injections.

**Figure 3.**
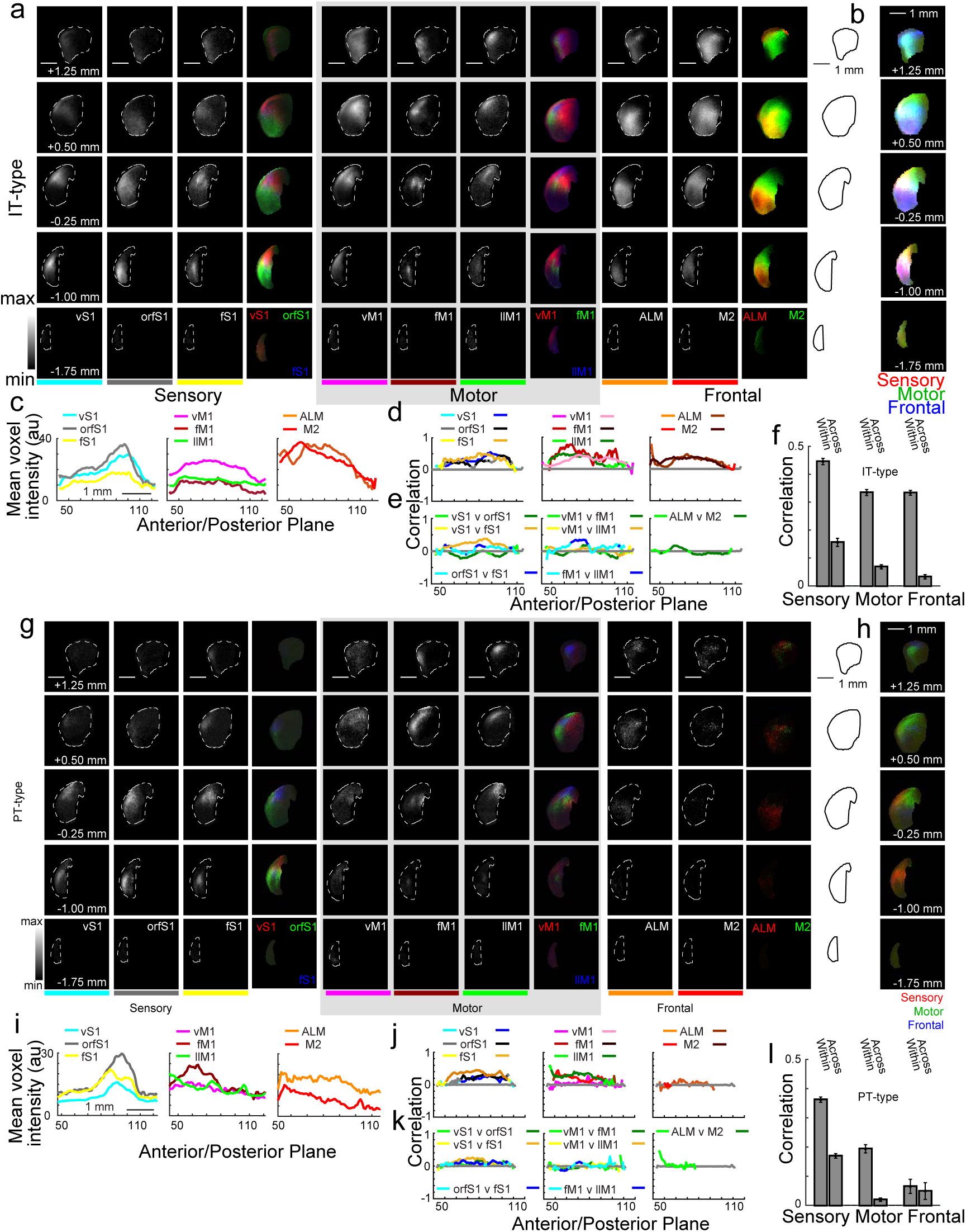
Topography of sensory, motor, and frontal corticostriatal projections from IT-type and PT-type neurons. (a)Images of average corticostriatal projections from IT-type neurons. Rows represent images at five coronal planes from anterior (+1.25 mm to bregma) to posterior (-1.75 mm to bregma). A dashed white line outlines ipsilateral striatum. Scale bar, 1 mm (top panel). Voxels are 50x50x50 μm. Columns represent the eight injection site clusters, organized into sensory (vS1, orfS1, and fS1), motor (vM1, fM1, llM1), and frontal (ALM and M2) modalities. Black and white images show average normalized projections for a given injection site cluster. For comparison, these are color coded and presented together at the right to show within-modality topography. For example, vS1 projections in red are generally more dorsal and orfS1 in green are generally more ventral. (b) Average normalized sensory (red), motor (green), and frontal (blue) projections are shown to illustrate topography across modalities. (c) Mean voxel intensity along the anterior/posterior axis of ipsilateral striatum. Scale bar, 1 mm; each plane is 50 μm. Each injection site cluster is color coded after Fig. 1. (d) Within cluster comparisons show high correlation for nearby injections in the same cluster. The sensory plot shows mean correlation for a given vS1 injection compared to other vS1 injections. Two colors are used (left, teal; right, blue) with the right-hand color indicating locations along the anterior/posterior axis where correlation coefficient is significantly different from shuffled data (p<0.001, rank sum test). Legend for all comparisons shows two colors for each injection site cluster (right color, significant differences). Similar comparison performed for all eight clusters. Comparisons made in planes for injections where both share >100 suprathreshold voxels. (e) Across cluster comparisons compare injections within the same modality. For sensory clusters, vS1 injections are compared to orfS1(green) and fS1 (yellow), and orfS1 injections are compared to fS1 (blue). Across cluster comparisons are also compared for motor (center) and frontal (right) injections. (f) Mean correlations within (vS1-vS1) and across (vS1-orfS1, etc.) ipsilateral corticostriatal projections from IT-type pyramidal neurons. Correlations within a given injection cluster are greater than correlations across functionally similar nearby clusters. (g-l) Images and analysis for PT-type projections, presented as for IT-type projections.

To assess corticostriatal somatotopy, quantitative comparisons were made between injections in the same injection cluster (Fig. 3d) or across injection sites of the same modality (Fig. 3e) using the methods described (Fig. 2). Comparison of correlation coefficients between injections within the same cluster (vS1 to other vS1 injections, Fig. 3d), showed these were always positively correlated. However, there was remarkably little correlation between injection sites across clusters of the same modality (Fig. 3e-f; Supplementary Fig. 3). ALM compared to the other frontal injection, M2, showed near-zero correlation, as did vM1-llM1, vS1-orfS1, and orfS1-fS1 comparisons. Where there was positive correlation observed in across-cluster comparisons, this was weaker than within-cluster comparisons. This suggested stereotypy in axonal projection patterns across mice. Contralateral corticostriatal projections (Supplementary Fig. 4) had grossly similar results with weaker overall correlations. Frontal areas, however, had particularly strong contralateral projections and similarly strong within-cluster correlation. This demonstrated that striatal targets of somatosensory and motor areas recapitulated some aspects of cortical topography.

This analysis was repeated for PT-type projections grouped into the same eight clusters by injection site location (Fig. 3g-l). There were general similarities, with frontal and motor projections targeting more anterior sites and sensory projections targeting more posterior ones. In contrast to IT-type projections, PT-type projections from frontal areas had fewer suprathreshold voxels and showed reduced mean voxel intensity compared to IT-type tracing from the same region (Fig. 3i). This reduction in intensity was consistent with smaller projections and less overlap between different injection sites. Thus, sensory injections were more segregated posteriorly in PT-type injections (red in Fig. 3h) compared to IT-type ones (purple in Fig. 3b). Comparisons for nearby injections in the same cluster (vS1 to vS1) had higher positive correlations than comparisons to injections in nearby clusters, such as vS1-orfS1 or vS1-fS1 (Fig. 3j-l; Supplementary Fig. 3). The correlations for all within and across group comparisons were summarized in Fig. 3l. Correlation scores were always higher for within than across group comparisons. Furthermore, PT-type projections have lower correlations than IT-type ones (Fig. 3f, l).

### Somatotopic specificity differs between IT-type and PT-type projections and between sensory and motor areas

Because these injections densely sampled sensory and motor areas, somatotopic specificity could be examined by comparing injections at a range of distances in the same or different cell types. Injection sites from different mice in the same location of the CCF are expected to share high correlation in their projections if connections in the rodent brain were stereotypical. Barrel cortex, for example, is sufficiently stereotyped that individual barrels are apparent in the Allen averaged registration template^16^. In contrast, microstimulation maps for movement show some inter-animal variability^22,23^. To examine the relationship between the distance between injection sites and their projections, the distance between injection site centers of mass was calculated for IT-type or PT-type injections in sensory and motor cortex. The correlation score in ipsilateral striatum was plotted against injection site offset (Fig. 4). For both sensory (blue) and motor injections (pink; Fig. 4b,d,f), the correlation score was fit with a linear regression (95% confidence interval shown). For IT-type projections, the peak correlation was higher for sensory cortical injections (∼0.6) than for motor cortex (∼0.4). The relationship dropped off more steeply in sensory areas (ANOCOVA, Group*X Value, p<0.0001). Collectively, these results suggest that sensory cortical areas show stronger topography than motor ones^22,23,26,28-30^. A similar relationship was apparent for PT-type projections, with higher correlations in nearby sensory injections than in motor areas (ANOCOVA, Group*X Value, p<0.0001). Peak correlation was stronger for IT-type than PT-type projections for both sensory and motor populations.

**Figure 4.**
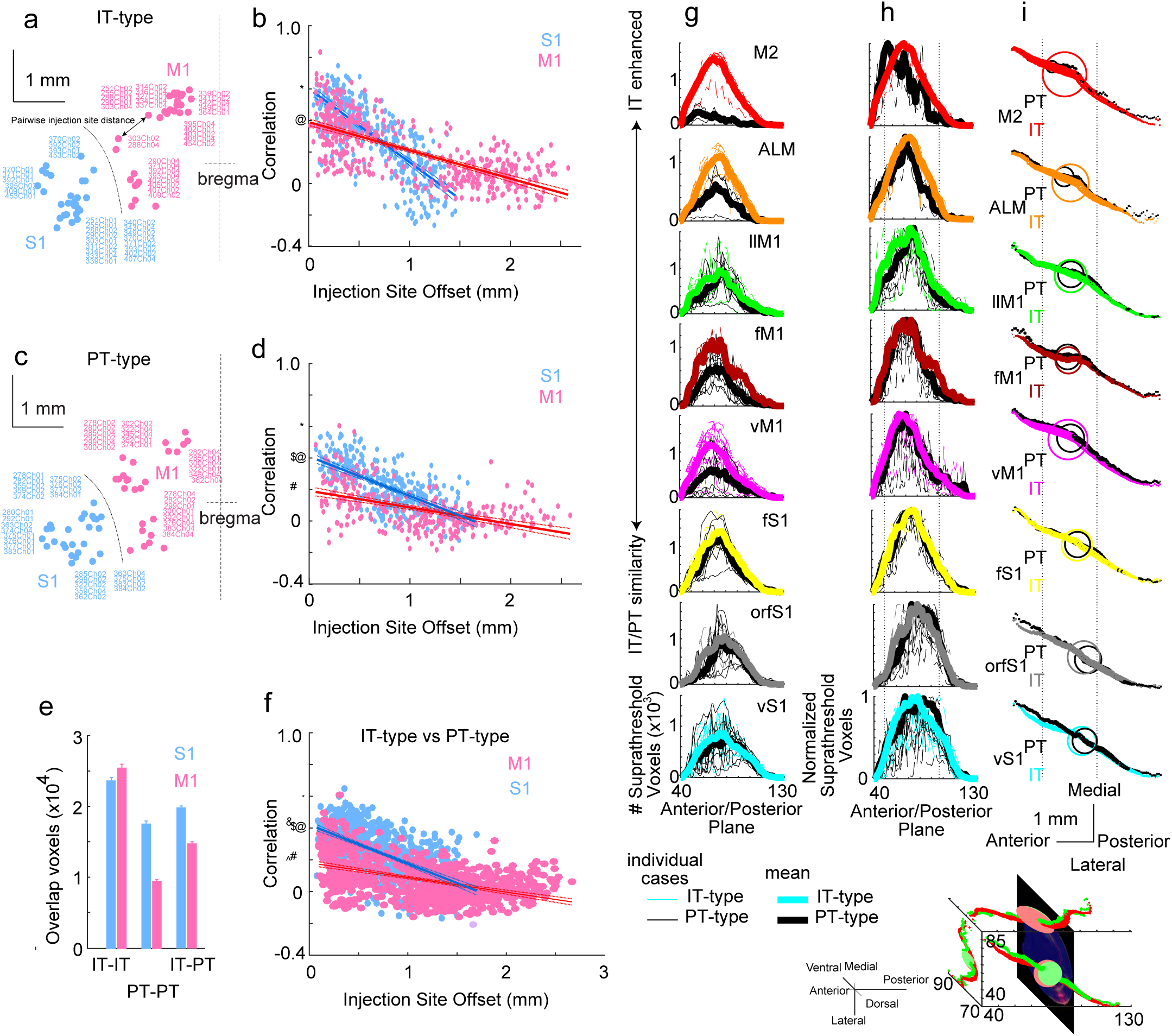
Somatotopic precision across the corticostriatal projectome compared across cortical areas and cell types. (a,b) Pairwise correlation between projections to ipsilateral striatum from IT-type (a-b) and PT-type (c-d) pyramidal neurons was determined and plotted against injection site offset in mm. (a, c) Dorsal view of injection sites in CCF coordinates. Midline and bregma indicated at right. Scale bars, 1 mm. All primary sensory (S1) injections are shown in blue and primary motor (M1) injections are shown in pink. Circles indicate injection site with injection number labeled. Double headed arrow indicates injection site offset distance for one pair of injections. (b, d, f) Correlation versus injection site offset for S1 and M1 injections. Solid line represents linear fit, with confidence interval plotted as dashed lines. Typographic marks indicate y-intercept across panels for comparison. (e) Mean number of overlapping voxels used to calculate correlations for IT-IT (b), PT-PT (d), and IT-PT (f) comparisons. (f) Correlation versus injection site offset for comparisons across IT-type and PT-type injections in S1 and M1. Here, each S1 IT-type injection is compared to each S1 PT-type injection but not to other IT-type injections. (g) The anterior/posterior location of suprathreshold voxels in ipsilateral striatum was quantified for all individual IT-type and PT-type injection cases. Individual cases are shown as thin dashed lines, while thicker lines represent the mean. IT-type projections are highlighted in color corresponding to their injection cluster (for example, vS1 is teal) while the corresponding PT-type projections from the same cluster are plotted in black on the same axes for comparison. Number of suprathreshold voxels is similar for vS1, orfS1, and fS1 injections. Suprathreshold voxels for IT-type projections from frontal areas ALM and M2 exceed those of PT-type projections. (h) Peak normalized distribution of both IT-type and PT-type projections are shown. These peak at similar points on the anterior/posterior axis. (i) To assess differences in targeting of IT-type and PT-type projections within the same injection site cluster, the center of mass of the voxels for the mean normalized injection pattern was calculated for each injection site cluster. The overall center of mass is shown as a large circle (red and green circles, example at left). The center of mass of each coronal plane is also plotted as a circle, and projections along the x-, y-, and z-axes are shown. The size of the circle is proportional to the summed normalized voxel intensity for a given plane. For the example projection at bottom, red (vS1 IT-type projection) and green (vS1 PT-type projection). The anterior/posterior projections for each injection cluster are shown above. The color code corresponds to the injection site cluster (teal for vS1), with PL56 injections shown in color and corresponding PT-type projections shown in black. Dotted line is shown for anterior/posterior alignment across injection clusters. Center of mass of vS1, orfS1, and fS1 (teal, gray, and gold, respectively) are posterior within the striatum, while frontal areas ALM and M2 (orange and red) are anterior. The overall center of mass of projections overlaps for IT- and PT-type cases, resulting in overlap of these markers.

The correlation of IT-type with PT-type injections near the same site was also studied. If these projections targeted different striatal regions, then both a reduction in the correlation as well as a reduction in the number of overlapping voxels were expected. However, the correlation versus distance relationship was similar to that of the within PT-type injection comparisons (Fig. 4f) while the number of overlapping voxels was intermediate to IT-IT and PT-PT comparisons (Fig. 4e). This was consistent with the center of mass of these injections falling in generally the same portions of striatum (Fig. 4g-i). Differences in these correlations could thus not be attributed to IT-type and PT-type neurons from the same cortical area targeting largely distinct striatal regions.

The departure from perfect correlation between projections from nearly overlapping injection sites could result from differences in the injection size (including number of infected cells and scatter at the injection site), inter-animal variability, or noise in image acquisition. Thus, whether different degrees of injection site scatter resulted in less correlation was tested. Injection site scatter was measured as the standard deviation for each infected soma from the injection site center of mass in a given injection. This was used to divide injections into two categories: those with scatter higher or lower than the mean. Correlation of ipsilateral striatal projections for low and high scatter groups was compared (Supplementary Fig. 5). Two populations were nearly indistinguishable, suggesting that injection size was not a major contributor to differences in correlations.

One model of corticostriatal organization suggests that striatal regions integrate input from multiple interconnected cortical areas^4^. This predicts that reciprocally connected regions of sensory and motor cortex would have elevated correlation in their striatal projections. Thus, pairwise comparisons between motor and sensory injections were examined. To assess the degree of corticocortical correlation, overlap of sensory axons in motor cortex (M1) injection sites (or motor axons in sensory cortex (S1) injection sites) was assessed. The M1 and S1 injection sites were defined in the CCF using coordinates that encompassed all labeled somata at the motor or sensory injection site, and included all voxels from pia to white matter. The correlation between a pair of M1 and S1 injections was then determined in this cortical volume, using the methods described in Fig. 2. Scatterplots compare the corticocortical correlation to the corticostriatal correlation for the same pair of injections (teal arrows, Fig. 5d-e). Each point represents the comparison of a single pair of injections. Red points specifically highlight comparisons between sensory and motor injections. Black points label pairwise comparisons between frontal areas and either motor (Fig. 5d) or sensory cortex (Fig. 5e). For IT-type projections, there was a positive relationship for striatal comparisons to M1 and S1 injection sites (Fig. 5d-e; R^2^=0.3640 for striatal correlation vs M1 injection site correlation; R^2^=0.3055 for S1 injection site correlation). In contrast, PT-type projections did not show this strong relationship (Fig. 5h-I and Supplementary Fig. 6; R^2^=0.0038 for striatal correlation vs M1 injection site correlation; R^2^=0.1219 for S1 injection site correlation). Because IT-type corticostriatal projections generally project to a greater area in striatum (Fig. 3 and 4), it is possible the increased co-correlation resulted from IT-type projection overlap in a focal region not innervated by PT-type neurons. Thus, the relationship between anterior/posterior subsets of the striatum with corticocortical connectivity (measured as before) was assessed by examining the co-correlation of cortical and striatal connectivity along 250 μm striatal segments. This revealed a long plateau of high correlation across the rostrocaudal extent of striatum (Fig. 5j) for IT-type but not PT-type projections. The enhanced co-correlation for IT-type projects did not result from a single focal region, but was spread across the extent of the corticostriatal projection. Thus, interconnected cortical areas shared projection targets in basal ganglia, but this relationship was stronger for the IT-type subset of corticostriatal projections.

**Figure 5.**
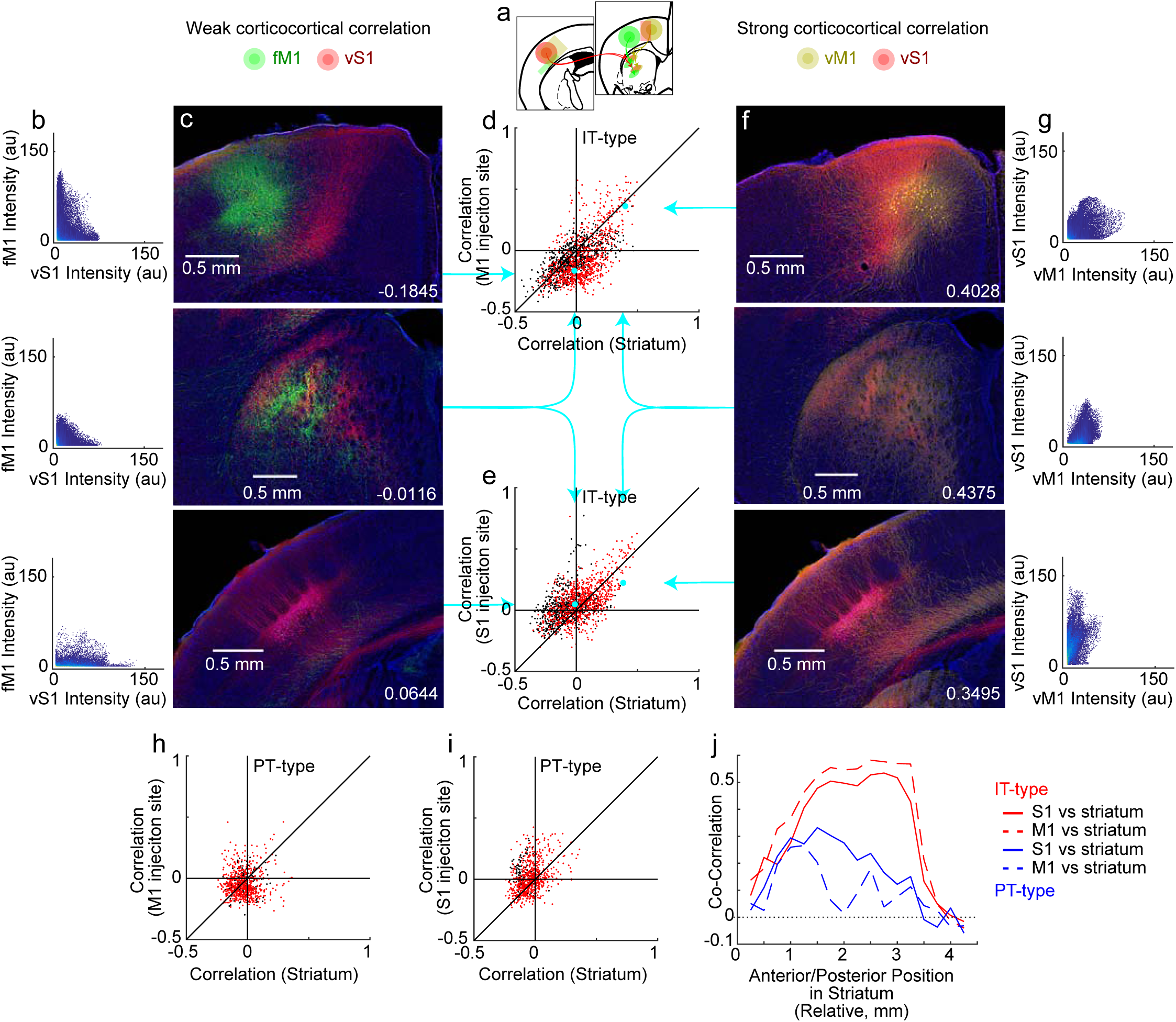
Corticostriatal projections map the organization of corticocortical connectivity in IT-type but not PT-type projections. (a)Sensory and motor cortex injections make reciprocal intracortical projections between somatotopically related areas. (b, c, f, g) IT-type injection examples shown contrast a pair of strongly connected cortical areas (red vS1 and yellow vM1 injections) with a non-somatotopically aligned area (green fM1 injection). vS1 axons (red) overlap poorly with fM1 neurons (green). These are poorly correlated in both injection sites (-0.1845 and 0.0644; b, c top and bottom) as well as the striatum (-0.0116; b, c middle). In contrast, vS1 axons (red) overlap well with vM1 neurons (yellow) and are strongly correlated in both injection sites (0.4028 and 0.3495; f, g top and bottom) as well as the striatum (0.4375; f, g middle). (d,e) Scatterplot of co-correlations of corticocortical connectivity (using injection site overlap) and corticostriatal connectivity for IT-type projections. Each individual point represents the corticostriatal correlations (x-axis) and injection site correlation (y-axis) for a single pair of injections with corticocortical correlation computed at either M1 (d) or S1 injection sites (e). Red points on the scatterplot compare sensory and motor injections. Black points add comparisons to frontal areas (M2 and ALM). Teal arrows and points indicate specific points corresponding to the example injections shown. (h,i) Scatterplot of co-correlations of corticocortical connectivity and corticostriatal connectivity for PT-type projections. (j) Co-correlations of corticocortical connectivity and corticostriatal connectivity are re-assessed, with corticostriatal correlations (y-axis) calculated using subsets of striatal voxels along the anterior/posterior axis in 250 μm segments (x-axis, in mm). Co-correlation is plotted for IT-type (red) and PT-type (blue) injections.

### Single IT-type and PT-type axons show similar gross targeting but differences in stereotypy and density of arborization

Mean projections were based on ∼600-900 neurons per injection (IT-type injections 906.9±71.7, PT-type injections 612.1±44.7, mean±sd). Examination of axonal arbors of single neurons shed light on how variable the projections of each population of pyramidal neurons might be. Single axons of IT- and PT-type cells in primary and secondary motor cortex were imaged and registered to the Allen Reference Atlas^31^. Although a limited number of total neurons were available, individual axons extended certain aspects of these findings. IT-type and PT-type neurons in the same area shared a similar topography, though IT-type arbors were more extensive and PT-type arbors were more focal (Fig. 6a-c). Comparison of multiple primary motor cortex (M1) projections confirmed that larger IT-type arbors have more overlap, while more focal PT-type projections were less likely to overlap. From the same area, PT-type axons innervated a subset of the region innervated by IT-type axons (Fig. 6d-f). The overall pattern of IT-type projections differed between M1 and M2 (Fig. 6g-i). M1 axonal projections targeted more discrete areas, with relatively nearby neurons maintaining somatotopic organization. In contrast, individual M2 axons projected more broadly within the striatum, resulting in considerable overlap and only a rough somatotopic organization. M2 projections were also stronger to contralateral striatum. Individual IT-type neurons in M1 and M2 showed considerable heterogeneity in terms of bilateral projections, with some neurons projecting axons primarily ipsilaterally, some contralaterally, and some bilaterally (cf. IT-type gold vs. red). Considerable variation between individual IT- and PT-type neurons suggested that further subclassification of these cell types is needed.

**Figure 6.**
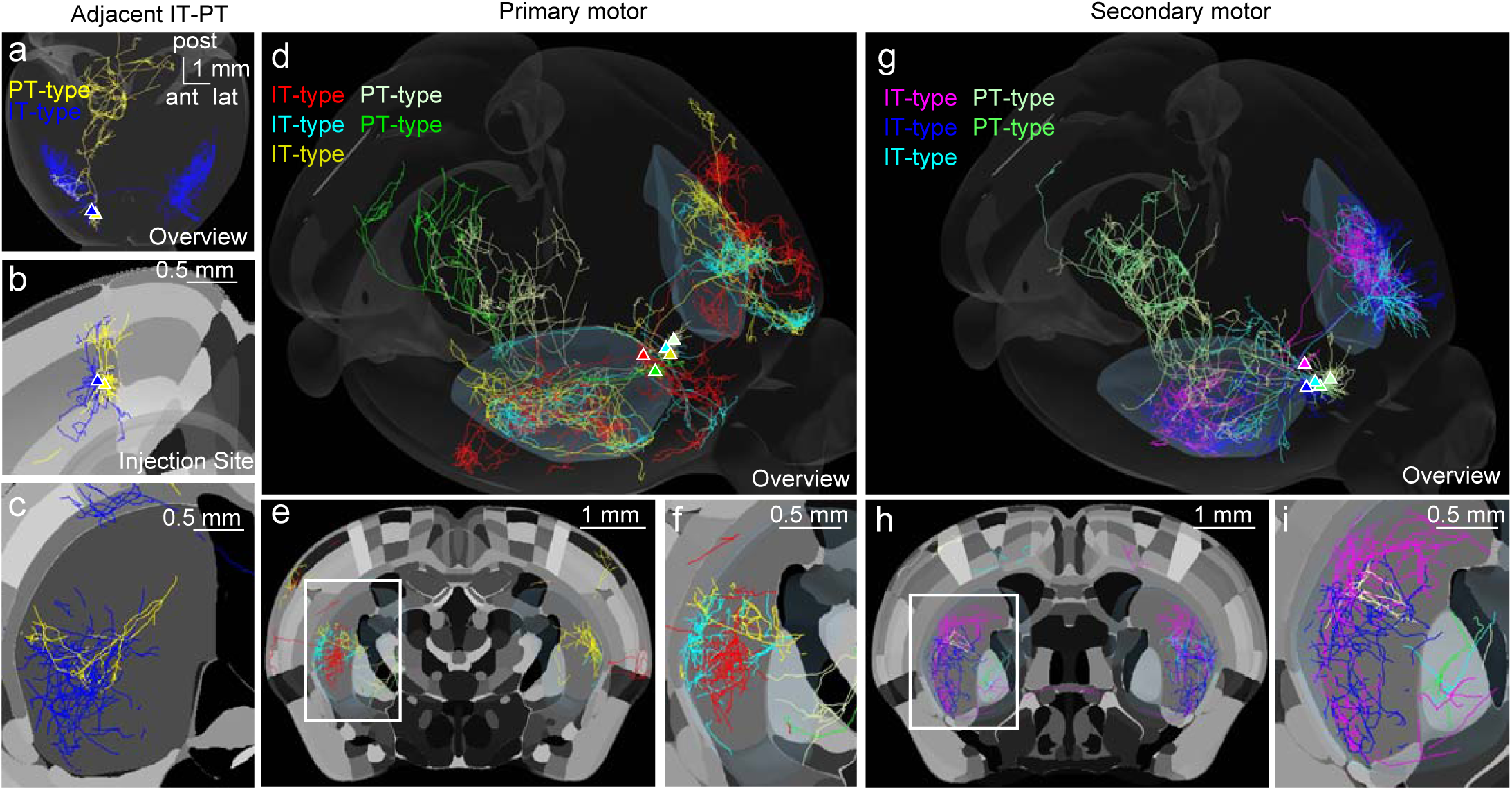
Single neuron reconstructions of IT-type and PT-type neurons in motor areas. (a)Reconstruction of the long-range axonal projections of adjacent PT-type (blue) and IT-type (gold) L5 pyramidal neurons^31^. Projections throughout the whole CNS anterior to medulla are shown in the reference atlas (CCF) coordinates, with annotated regions shown in gray. (b) Somata are in adjacent in primary motor cortex (M1). (c) Corticostriatal projections show similar general topography with differences in arbor size and density. (d) Five adjacent IT-type (red, teal, and gold) and PT-type (green and white) M1 neurons. (e,f) Corticostriatal projections show topography of IT-type and PT-type projections, as well as differences in density, terminal field size, and asymmetry of projections to contralateral striatum. White box in (e) magnified in (f). (g-i) Five adjacent IT-type (purple, teal, and blue) and PT-type (green and off-white) secondary motor cortex (M2) neurons.

### Striatum is loosely organized in somatotopic areas

IT-type and PT-type projection correlations were used to construct hierarchical relationships between cortical injection sites based on the projections to various brain regions. Pairwise correlation scores for IT-type outputs to ipsilateral striatum were used to construct a dendrogram using Euclidean distance between correlations as the distance measure. Generally, nearby injection sites showed the greatest affinity (Fig. 7a-c). At higher hierarchical levels, most fS1 and vS1 injections clustered together. Motor injections in vM1, fM1, llM1, and M2 also clustered together. Unexpectedly, orfS1 clustered with ALM, suggesting an affinity between lateral sensory and frontal areas in their projections to ipsilateral striatum. Of interest, this affinity also recurred in a similar analysis of corticocortical correlations (Supplementary Fig. 7). In contrast to the IT-type results, using the same methodology to examine PT-type corticostriatal outputs, sensory inputs clustered together, separately from motor and frontal inputs (Fig. 7d-f).

**Figure 7.**
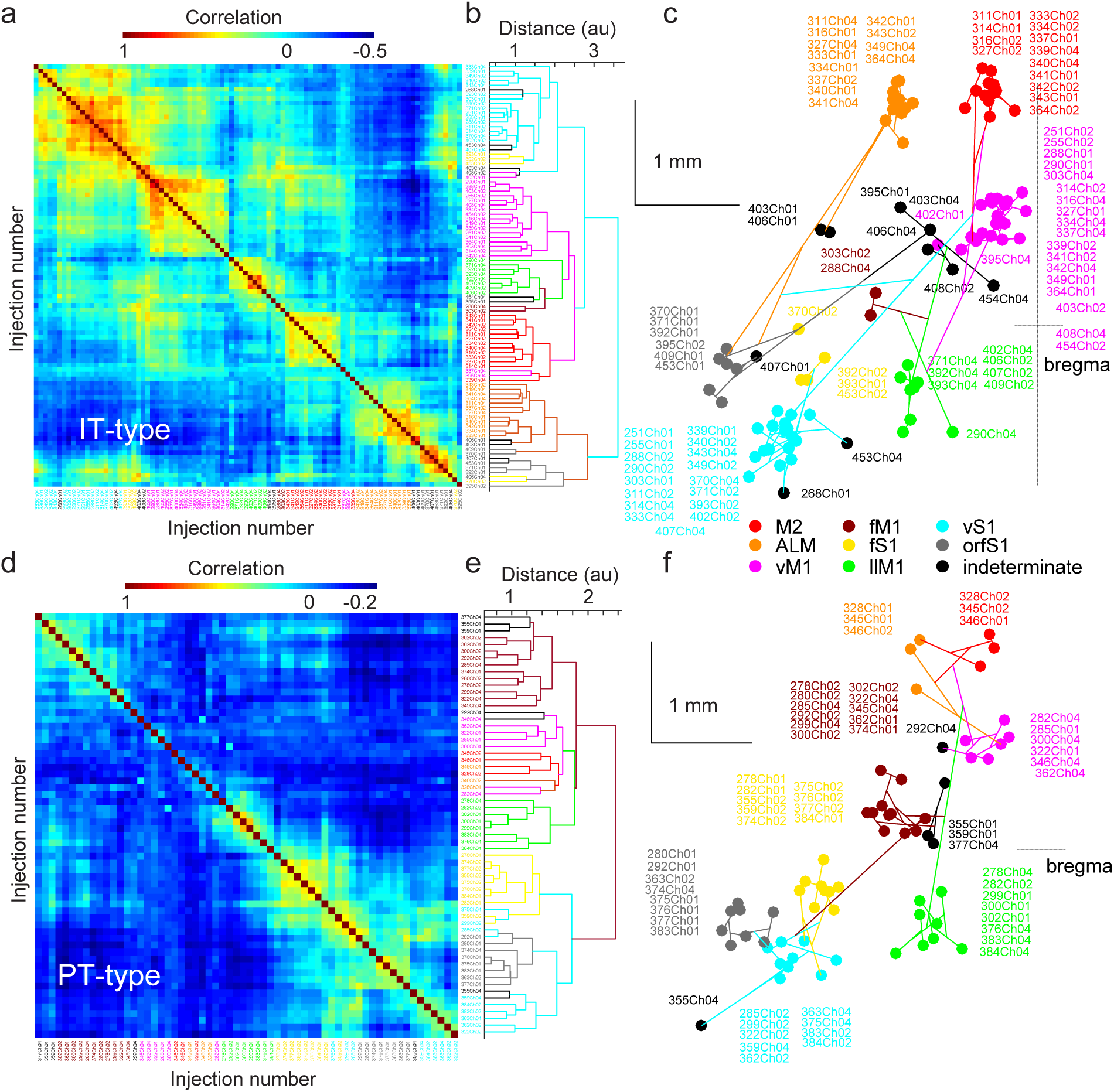
Hierarchical clustering of IT-type and PT-type corticostriatal projections. (a)Pairwise correlation scores for all IT-type projections studied (N=92). High correlation, red (with perfect correlation along the main diagonal). Negative correlation, blue. (b) Injections were hierarchically clustered using correlation score as the distance measure. Individual injections at the tips of the dendrogram were color-coded according to the injection site location cluster to which they were assigned. (c) Using a dorsal view of the brain (with bregma marked at right; scale bars, 1 mm), the dendrogram from (b) was plotted using the center of mass of the injection site as the point for the tip of the tree. (d-f) Pairwise correlation scores and dendrograms for all PT-type projections studied (N=62), plotted as for IT-type projections.

Differences in input contribute to differences in striatal function. Since corticostriatal inputs form a major excitatory input, differences in sensory, motor, and frontal corticostriatal projections could identify functionally distinct striatal regions. Average normalized projection patterns were determined from eight injection sites for two mouse lines. The normalized projection strength was used to assign ipsilateral striatal voxels into clusters using *k*-means clustering. Five clusters were found based on the peak silhouette value. These were presented in coronal section for the ipsilateral striatum using five colors (Fig. 8a). The fraction of output to each of the clusters is shown for IT-type and PT-type projections (Fig. 8b-c). One cluster (blue) covered the anterior, medial, and posterior edges of the striatum, which were predominantly regions receiving poor output from sensorimotor cortex. The dorsolateral sector included (a) an anterior core region (green) that received substantial M2 and primary motor output, (b) an anterior dorsolateral region (olive) that received strong motor output and some sensory output, and (c) a posterior dorsolateral region (red) that received strong sensory output and some motor output. The ventral and posterior domain received input from ALM and orfS1. This analysis was repeated for IT-type projections alone and PT-type projections alone (Supplementary Fig. 8). Clustering based on IT-type input alone resulted in 4 clusters, with the anterior and posterior dorsolateral regions that were separable based on both projections combined into a single cluster when PT-type data was excluded. This shift highlighted a difference in the IT-type and PT-type projections: the primary motor projections favored the anterior (olive) dorsolateral cluster, while the primary sensory projections favored the posterior (red) dorsolateral cluster. This difference was more pronounced for PT-type than for IT-type. Thus, differences in PT-type projections identified putative functionally distinct regions of striatum. That these regions were divided by PT-type sensory and motor outputs is also consistent with the earlier dendrogram (Fig. 7). The clustering of IT-type outputs to contralateral striatum was similar to ipsilateral striatum, but not as well-defined. Three clusters were sufficient to describe contralateral projections (Supplementary Fig. 8). Consistent with this, the overall correlation coefficients were reduced for these projections (Supplementary Fig. 4). This implied a reduction in the topographic specificity of contralateral striatal projections.

**Figure 8.**
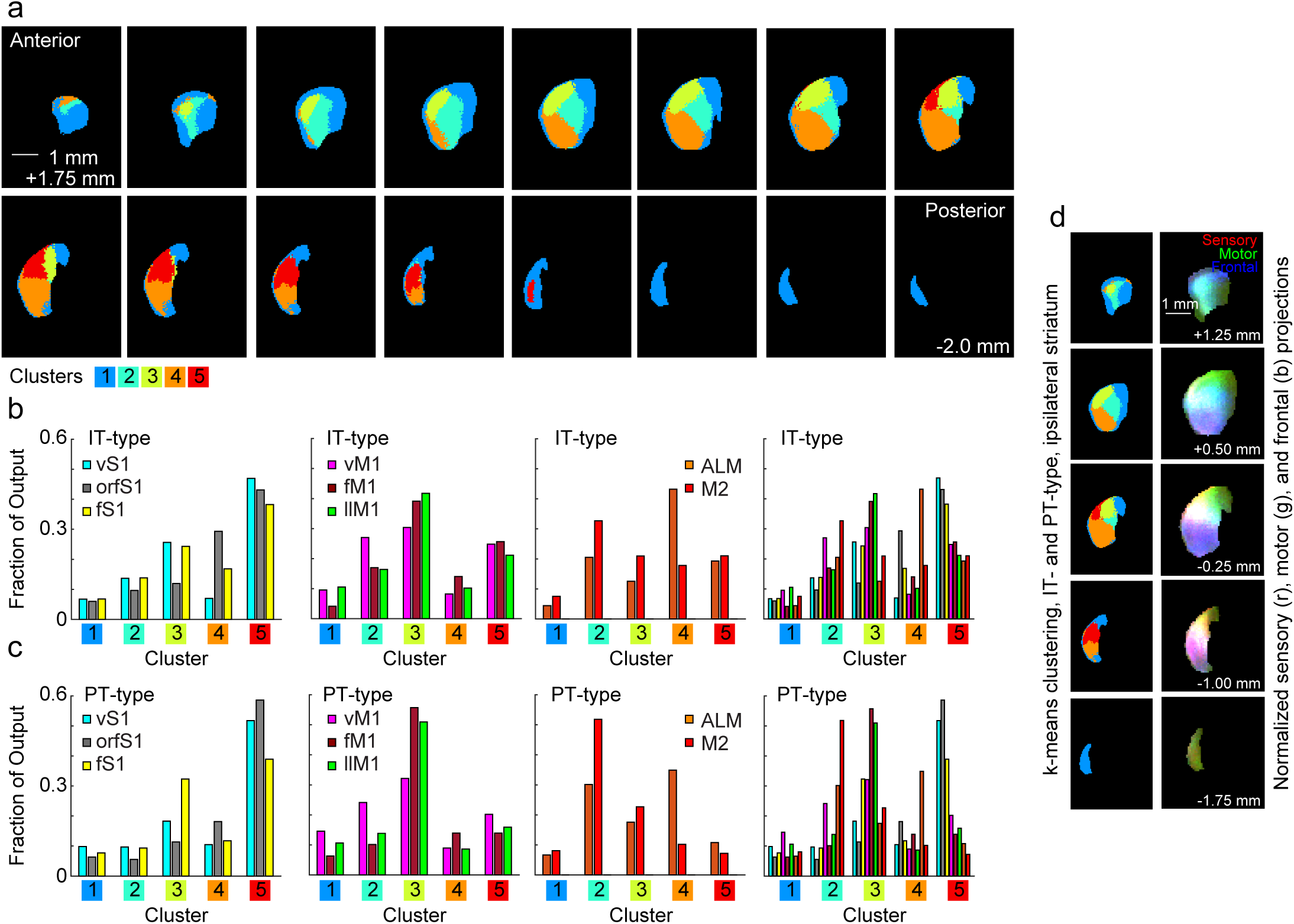
Major divisions of ipsilateral striatum based on sensory and motor cortical projections. (a)*k*-means clustering of striatal pixels based on mean normalized fluorescence intensity from each of the eight injection site clusters for both IT-type and PT-type pyramidal neurons. The striatal clusters are illustrated as five colors (legend, at bottom) in evenly spaced planes every 0.25 mm from anterior (top left) to posterior (bottom right). Scale bar, 1 mm. (b) The fraction of the output from each of the eight injection site clusters to a given striatal division from IT-type projections. Graphs are divided into sensory (left), motor (center), and frontal (right), with all areas together at far right. (c) The fraction of the output from each of the eight injection site clusters to a given striatal division from PT-type projections, presented as in (b). (d) A comparison of the striatal divisions based on *k*-means clustering (left) to the pattern of normalized sensory (red), motor (green), and frontal projections (blue), presented as an RGB image (right).

## Discussion

### Corticocortical connectivity predicts corticostriatal convergence of output from specific cell types

How do the corticostriatal projections of PT-type and IT-type neurons differ? These results show that corticostriatal projections of interconnected cortical areas replicate their corticocortical connectivity by projecting to shared targets in the striatum^3,4^ (Fig. 5). However, this model did not distinguish between cell type specific projections. Discriminating between IT-type afferents and PT-type collaterals revealed this model best describes IT-type projections. Differences in the corticostriatal topography of projections for specific cell types within a cortical region had not previously been predicted. The basis for this difference is not that the center of mass of these projections differs (Fig. 4). Instead, within nearby cortical sites, there is greater heterogeneity in the PT-type projection between animals, as well as between axonal projections of single cells, as seen in MouseLight (Fig. 6). Thus, PT-type output is more focal, but less stereotyped in its targeting, as evidenced by both population and single axon data. These differences have been difficult to appreciate with conventional tracing techniques, though overlapping projections in subcortical targets including thalamus have been effective as a measure of somatotopic alignment between cortical sites^1^. Fine afferents may be missed in the corticostriatal projection in the Golgi method^32^ and tracers do not distinguish between cell types^2,24^. Thus, cell type-specific lines are advantageous for anatomical tracing since the long-range projections of different cell types are organized differently^15^.

The relative importance of IT-type and PT-type corticostriatal collaterals is unclear. Both cell types are significant in rodents, as seen here. PT-type collaterals are also present in primates^33^, but are less prominent^13,34^. These neuronal subtypes receive distinct inputs^10^ and convey different classes of information to descending circuits^13^. Thus, these differences may contribute to functional specialization within the striatum. These quantitative measures would be difficult to achieve with lower resolution alignment (>100 μm voxels) or the scoring of axons as present or absent (reducing the bit depth of images), which may limit similar studies^2,24,35^. Inclusion of other subtypes of projections, such as L2/3 pyramidal neurons or thalamic inputs^35^, or further subdividing IT-type neurons (as is possible with MouseLight) may reveal more nuanced structure within the corticostriatal projection.

### Differences in somatosensory and motor topography

The difference in correlation between nearby primary motor and somatosensory projections is remarkable. In comparing IT-type injections in S1 and M1, the highest correlations are found for nearby injections in S1 (Fig. 4). The higher correlation with steeper reduction as injection sites shifted apart is consistent with a greater topographic specificity in primary somatosensory areas. This is paralleled by functional data, where specific areas of S1 are highly specific for certain body regions such as barrel cortex, where individual barrels are specific for a single whisker^25^. In contrast, microstimulation data suggests that motor representations, while topographic, are also generally intermingled^22,23,26,29,30,36,37^. The basis of these somatotopic differences may derive from the fact that somatosensory cortical areas have a clearly defined input for a given cortical column, such as the primary thalamocortical afferent to layer 4, representing touch of a single finger or whisker^38^. In contrast, primary motor areas have less spatially restricted thalamic^39^ and cortical^40^ inputs. The neurons in these areas may represent a more diverse range of phenomena^41^, ranging from muscles^42,43^ to movements^44^ and behaviors^45^, where body representation alone is not sufficient. It is worth noting that the decrease of correlation with injection site offset is relatively linear instead of stepwise, though smaller steps in the noise are possible. This is consistent with a gradual shift in somatotopic representation of body regions in striatum instead of discrete segments dedicated exclusively to a single region^2^.

This relationship is also true between sensory and motor injections labeling PT-type neurons, but the overall level of correlation is lower. This was unexpected, as these projections, as collaterals of output targeting subcortical targets, were expected to be more precise. The enhanced correlation of IT-type neurons is not due to targeting of a specialized IT-specific striatal region or a substantial offset in the projection zones of the two cell types, as the center of mass of PT- and IT-type projection is similar across the anterior/posterior extent of the striatum (Fig. 4g-i). Instead, quantification of PT-type collaterals showed that these projections have fewer suprathreshold voxels and thus are more spatially limited (Fig. 3-4). Individual axon reconstructions, such as MouseLight data, show that striatal axons of IT neurons are more highly branched than those of PT neurons^9^. Therefore, individual PT-terminals are more focal (Fig. 6). But they also show less spatial overlap and higher variability within an injection site (Fig. 4) and between nearby cells (Fig. 6). This correlation is not simply due to a reduction in the volume of overlap, as comparisons between PT- and IT-type injections in nearby sites showed an increase in overlap volume, but relatively low correlations comparable to PT-PT correlations for the same injection site offset (Fig. 4). Thus, peak correlation is not simply driven by overlap volume.

Although there is strong evidence from primates^4^ and rodents^46^ for convergence of corticostriatal afferents from associated cortical areas, some data^47^ suggests S1 and M1 projections are largely non-overlapping. This result may differ from those presented here if the somatotopic alignment of the two sites is imprecise (Fig. 4 and 5). The dual channel recordings presented here (Supplementary Fig. 2) show synaptic convergence of S1 and M1 outputs for all SPNs recorded, demonstrating that integration of somatotopically aligned sensory and motor signals is a relatively frequent characteristic of striatal neurons.

Contralateral corticostriatal projections of IT-type neurons show reduced correlations compared to ipsilateral axons (Supplementary Fig. 4). Thus, the precision of axonal targeting varies across different collaterals of the same cell type. Since it would be possible to use the same molecular and activity-dependent cues to achieve the same precision in ipsi- and contralateral connections, it will be interesting to learn the functional import of generating a contralateral projection with less spatial precision than the ipsilateral one. On the one hand, longer-range contralateral projections might lose some topographic precision, but how does the animal benefit from a less precise contralateral projection? Such inputs would seemingly degrade the precision of input to contralateral SPNs.

Notably, overall projection density differs across IT-type and PT-type neurons moving from frontal to motor and sensory areas (Fig. 4). In IT-type injections, frontal projections provided the densest striatal afferents (Fig. 3). In contrast, for PT-type injections, frontal injections were by contrast the weakest (Fig. 3 and 4). Thus, PT-type projections had a higher relative density of projections from sensory areas. This difference is useful in subdividing the striatum into sectors, where including both PT- and IT-type projection data helps differentiate anterior and posterior dorsolateral striatal areas specialized for motor and sensory input respectively (Fig. 8, clusters 3 and 5) which merge when IT-type only output is considered (Supplementary Fig. 8a).

Several sources may limit the stereotypy of corticostriatal projections. Relatively compact versus scattered injection sites did not show a large variation in corticostriatal correlations, suggesting that injection site size did not play a major role in variation between injections. Thus, animal-to-animal variation instead of injection variability may play a larger role in limiting the peak correlation. Other limitations include the spatial resolution of the alignment (∼50-70 μm) and voxel size, which could reduce correlations by spatial averaging. Using higher resolution aligned images (10x10x10 μm voxels) did not alter the linkages between injection sites (data not shown). That peak correlations are close to 0.6 suggests that animal-to-animal variation sets an upper limit on comparisons across brains. That peak correlations are not closer to 1.0 quantified the substantial inter-case variability and underscores the relevance of studying injections across cases in different animals instead of using a single injection case to assess typical projection targets in striatum.

### Affinity of orofacial sensory and motor areas

Although frontal areas, such as ALM and M2, might be organized differently than sensory and primary motor cortex, it was interesting that IT-type projections from lateral regions of frontal cortex (ALM) projected to striatum similarly to those originating from orofacial regions of S1 anterior and lateral to barrel cortex. Of note, the corticocortical collaterals of ALM also projected posteriorly towards lateral regions of motor and somatosensory cortex. This was reciprocated by projections from orfS1 to ALM. Thus, ALM’s corticocortical connectivity suggested a basis for corticostriatal overlap with orfS1 projections. Coincidentally, ALM has been identified as a low-threshold region for evoking tongue movement in rodents^22,48,49^. Based on this connectivity pattern, ALM and orfS1 are connected in a manner reminiscent of primary motor and sensory regions. ALM has also been implicated in more traditional frontal cortex functions such as motor planning in mice^50,51^.

### Limitations of anatomical methods

A comprehensive study of differences in cortical cell type (IT- or PT-type) output to distinct striatal populations was not possible from all cortical areas to the range of striatal neuron populations, including direct and indirect SPNs as well as striatal interneuron populations. Targeting of afferents to distinct striatal compartments such as patch and matrix may also differ between different cortical areas, though average sensorimotor populations target both patch and matrix neurons in similar proportions^52^. Afferents from both IT- and PT-type cells form connections to direct and indirect pathway striatal projection neurons^53^. It not yet possible to evaluate, however, whether there is a bias in targeting from either PT- or IT-type output, as has been proposed^54,55^, because of quantitative limitations in circuit mapping methods. Retrograde tracing with transgenic rabies suggests that sensory and motor inputs preferentially excite direct and indirect pathway SPNs respectively^56^, suggesting specific postsynaptic targeting of afferents is possible in striatum. Physiological data (Supplementary Fig. 2) shows that motor and sensory corticostriatal afferents converged on single SPNs (Supplementary Fig. 2), but did not quantitatively distinguish between the cell types targeted.

Layer-specific Cre-driver lines such as Tlx3_PL56 and Sim1_KJ18 lines may not collectively label all L5 pyramidal neurons. For example, in the L5 mouse line Rbp4_KL100, some IT-type and PT-type neurons are labeled, but the overall labelling density leaves many cells of both classes unlabeled^17^. The density of PT-type and IT-type neurons in Sim1_KJ18 and Tlx3_PL56 lines varies over cortical areas, which suggests that some neurons may be missed in different regions. There may be underappreciated heterogeneity within these two L5 populations, such as different subtypes of IT neurons for different targets^57^. Furthermore, in ALM injections of frontal cortex in Sim1_KJ18 mice, some contralateral axonal projections are present. In other areas, such as midline cortical areas where lamination is less pronounced, transgenic reporters for these lines suggest changes in Cre expression, resulting in reduced Tlx3_PL56 and Sim1_KJ18 labeling^17^. Thus, use of transgenic approaches to target specific cell types is limited to the brain regions where these cell types are well-characterized.

## Conclusion

The corticostriatal projection formed by two populations of L5 pyramidal neurons conveys distinct functional information with distinct striatal targeting. IT-type neurons in sensory and motor areas target somatotopically organized domains of striatum and also overlap substantially with other cortical areas with which they are reciprocally connected. PT-type neurons, in contrast, show less overlap with reciprocally connected cortical areas. This difference suggests that the measured degree of topographic organization depends in part on the cell type considered. As these cell types convey distinct information to striatum, it remains to be determined what purpose this differential targeting serves.

## Acknowledgements

We thank David Robbe, Taehyeon Kim, Sandra Okoro, YingXin Zhang-Hooks, and Peter Strick for insightful comments on the figures and manuscript. We thank Jack Glaser, Paul Angstman, Nate O’Connor, Sue Tappan, Mike Fay and Scott Gerfen at MBF Bioscience for the collaboration to develop and use the BrainMaker software. This work was supported by a NARSAD Young Investigator Award to BMH and by NINDS/NIH (R01 NS103993).

## Author contributions

BMH, AEP, BSE, and CRG wrote analysis software needed to quantify the data. BE developed the BrainMaker software at MBF Bioscience to align the whole brain. RP performed all anatomical work for sectioning, immunostaining, and imaging. MF quantified soma locations for all injections. JJC performed all recordings in striatum for the dual channel photostimulation experiment. JW and JC produced single axon reconstructions with the MouseLight project at Janelia Research Campus. BMH and CRG conceived of the project, analyzed the data, and wrote the paper with contributions from all authors.

## Online Methods

### Injections

All breeding, surgical, and experimental procedures conformed to National Institutes of Health guidelines for mice and were approved by the Institutional Animal Care and Use Committees of University of Pittsburgh and Janelia Research Campus. Mice from four GENSAT BAC Cre-recombinase driver lines (Sepw1_NP39, N=7; Tlx3_PL56, N=33; Sim1_KJ18, N=22; and Ntsr1_GN220, N=5)^17^ were used to trace the projections of four populations of cortical pyramidal neurons. Mice of both sexes were injected at postnatal day P37.0±1.7 (mean ± se) and sacrificed after 2-3 weeks of expression. Stereotaxic injections were performed as previously described^40^, with all injections in the same hemisphere. Injection sites covered a range of somatotopic locations in primary somatosensory cortex and corresponding areas of motor and frontal cortex^22-24^. 30 nL per injection site of AAV-flex-XFPs were injected using a custom positive displacement injector via a pulled borosilicate glass micropipette. The generic AAV-flex-XFP refers to several tracing viruses used, including AAV2/1-CAG-flex-EGFP, AAV2/1-CAG-flex-tdTomato, and the GFP- and mRuby2-based spaghetti monster fluorescent proteins (smFPs) smFP-FLAG, smFP-Myc, smFP-V5, smFP-HA, Ruby2-FLAG, and Ruby2- OLLAS (Table 1)^18^. Injections were made into cortex (at 300-1100 μm depth). For injections into L5 and L6, virus was injected at two depths. Laminar specificity was achieved by Cre-recombinase instead of injection depth. Typically, three sites were injected per mouse. In some cases, fewer channels were quantified if expression was not usable in a given channel due to weak expression or marked spread of the virus away from the injection site.

### Histology, staining, and imaging

Mice were transcardially perfused with 4% paraformaldehyde in phosphate-buffered saline and postfixed overnight. Brains were then transferred to 20% sucrose in PBS for storage. Brains were sectioned at 80 μm and signal was immunoamplified. 1:100 dilution of Neurotrace Blue was used as a structural marker^19^. Sections were then imaged using Neurolucida (v2017, MBF Bioscience, Williston, VT) on a Zeiss Axioimager (Zeiss, Oberkochen, Germany) equipped with 10x objective, Ludl motorized stage and a Hamamatsu Orca Flash 4.0 camera (Hamamatsu, Hamamatsu City, Japan). Each section was comprised of an average of ∼100-200 image stacks collected in 10 μm steps. A single 3D image was first generated then a deeper field-of-view was achieved by collapsing images to a single plane using a DeepFocus algorithm ^17,19^ (**Supplementary Fig. 1b-e**) implemented in Neurolucida. Original images are available at: http://gerfenc.biolucida.net/link?l=Jl1tV7

### Whole brain reconstruction, image annotation, and registration

Tiled images were aligned to a standard coordinate system using BrainMaker software (MBF Bioscience, Williston, VT). Resulting serially-reconstructed brains contained 10 μm isotropic voxels (782x1086x1242) and were registered to the annotated Allen Mouse Common Coordinate Framework (CCF), Version 3 http://connectivity.brain-map.org)^14-16^. All brains were registered to this framework using Neurotrace Blue images as the structural marker and a two-stage registration process. The first stage constructed an average reference space that provides a representation of the average appearance of brains that have undergone histological sectioning, mounting, and staining specific to this study and in the same image modality (i.e. Neurotrace Blue). Registration of individual brains to this average reference space was found to be more robust than direct multimodal registration to the Allen CCF reference image.

The average reference image was constructed from 78 individual 3D brains in a manner similar to the Allen CCF, which incorporates 1675 individual brains with cytoarchitecture visualized with 2-photon auto-fluorescence^16^. In this study, the counterstain (Neurotrace Blue) channel for each individual brain was registered to a reference template, initialized as one of the individual brains resampled with a uniform voxel spacing. Multiple resolution registration optimized the 12 parameters of a 3D affine transform to minimize a normalized correlation metric between each brain and the template image. The reference template was then updated by resampling all individual brains with their respective affine transforms and computing a voxel-wise weighted average. Voxels that received a small number of contributions were discarded to correct for some tissue damage present in individual brains. A second pass registered each individual brain to the new template, updating the individual transforms. This process repeated until the template image stabilized.

The second stage involved registering the average reference image to the Allen CCF. 300 unique landmark points were identified in the average reference image and corresponding points in the Allen CCF 2-photon reference image. The positions of the landmark correspondences were used to construct a nonlinear transform that models deformation of a uniform mesh grid with B-splines. This transform was used to resample the Allen CCF annotation volume in the average reference image using nearest neighbor interpolation. The result, an average reference image and its spatially aligned annotation volume, constitutes the average reference atlas. The counterstain channel of individual brains in this study were registered with the average reference space by adjusting parameters of a 3D affine and 3D nonlinear B-spline transform to minimize a normalized correlation metric. Some but not all individual brains contributed to the average reference space. Measurement of alignment precision showed this was accurate to ∼50-70 μm (Supplementary Fig. 1l-y). Comparable studies use alignment methodologies with less precision (∼100 μm), larger voxels (100-150 μm per side)^35^ or images reduced from 8-bit to 2-bit (“dense/strong”, “moderate”, “diffuse/light”, etc.)^2,24^.

The recovered transform was used to map the locations of fluorescence and cell soma locations detected on fluorescent tracer channels. For quantification of injection site location, tiled images were imported into Neurolucida software (MBF Bioscience, Williston, VT) and soma locations were annotated using automated object detection with manual supervision. Nearest neighbor interpolation of the average reference space volume at the mapped positions provided the anatomical region assignment for each cell. Coordinates of the CCF for structures of interest (such as striatum) were used to identify voxels for quantification. These were divided into left and right hemispheres to distinguish between structures ipsilateral and contralateral to the injection site.

## Data analysis

Aligned brain images were downsampled to 50 μm isotropic voxels (156x217x248) using custom routines in FIJI software^58^. The annotated Allen Mouse CCF was also used at 10 μm and downsampled to 50 μm. The annotation was used to assign voxels to a given brain region (ipsilateral or contralateral striatum, for example). Both 10 μm and 50 μm images were converted from tifs into .mat files in Matlab (Mathworks, Natick, MA) for analysis with custom routines. Soma locations were similarly imported to Matlab.

### Life Sciences Reporting Summary

Further information on experimental design is available in the Life Science Reporting Summary.

### Data availability statement

The data that support the findings of this study are available from the corresponding authors upon request. Aligned images in 10 and 50 μm voxels for all brains, cell soma locations, and the corresponding masks used to identify brain regions (striatum, for example) are available on request. Custom Matlab code for data analysis is available on request. Original images of whole are freely available online at: http://gerfenc.biolucida.net/link?I=JI1tV7.

